# scFlow: A Scalable and Reproducible Analysis Pipeline for Single-Cell RNA Sequencing Data

**DOI:** 10.1101/2021.08.16.456499

**Authors:** Combiz Khozoie, Nurun Fancy, Mahdi M. Marjaneh, Alan E. Murphy, Paul M. Matthews, Nathan Skene

## Abstract

Advances in single-cell RNA-sequencing technology over the last decade have enabled exponential increases in throughput: datasets with over a million cells are becoming commonplace. The burgeoning scale of data generation, combined with the proliferation of alternative analysis methods, led us to develop the scFlow toolkit and the nf-core/scflow pipeline for reproducible, efficient, and scalable analyses of single-cell and single-nuclei RNA-sequencing data. The scFlow toolkit provides a higher level of abstraction on top of popular single-cell packages within an R ecosystem, while the nf-core/scflow Nextflow pipeline is built within the nf-core framework to enable compute infrastructure-independent deployment across all institutions and research facilities. Here we present our flexible pipeline, which leverages the advantages of containerization and the potential of Cloud computing for easy orchestration and scaling of the analysis of large case/control datasets by even non-expert users. We demonstrate the functionality of the analysis pipeline from sparse-matrix quality control through to insight discovery with examples of analysis of four recently published public datasets and describe the extensibility of scFlow as a modular, open-source tool for single-cell and single nuclei bioinformatic analyses.

## Introduction

Single-cell RNA sequencing (scRNA-seq) has enabled transcriptomic profiling at single-cell resolution, providing unprecedented insight into gene expression within cell populations (Shema et al., 2019). However, a satisfactory framework for standardized, computationally efficient analyses of scRNA-seq (or snRNA-seq) data has not been available to date. Lack of full community agreement on quality measures and standards for quality control, typically large analytical batch effects and multiple parameter optimisations necessary in current tools have confounded reproducibility of results. Moreover, the burgeoning scale of scRNA-seq datasets made possible by technological advances including integrated fluidic circuits, nanodroplets, and *in situ* barcoding, has led to a concomitant increase in computational demands for individual dataset, underscoring a need for efficient scaling, particularly with recognition of the value of meta-analyses (Aldridge and Teichmann, 2020). A comprehensive solution to these challenges has not been provided (Eisenstein, 2020). Nonetheless, the benefits of reproducible computational practices in the life sciences are clear and a source of extensive discourse in the literature (Baker, 2016; Perkel, 2020). The demand from governments, funders, and publishers for FAIR (findable, accessible, interoperable, and reusable) standards in data-driven sciences is highly pertinent to scRNA-seq analyses (Sansone et al., 2019). Better realisation of these goals for scRNA-seq can be promoted by standardisation of core elements in analysis pipelines to enable common approaches to annotating data for quality and its characterisation. Common metrics and a scalable analytical framework would better enable the integration, re-use, and repurposing of published datasets within and across diseases to drive novel discoveries (Grüning et al., 2018). Challenges toward the development of such a pipeline include the deluge of computational techniques for key analytical steps (Heiser and Lau, 2020), interoperability challenges between analytical tools (Tekman et al., 2020), the extensiveness of complete parameter specifications (Raimundo et al., 2020), the iterative nature of hyperparameter optimisation (Menon, 2019), the complexity of software dependencies for end-to-end analyses (Gruening et al., 2018), and the need for flexibility to handle complex experimental designs (Luecken and Theis, 2019).

To this end, we have developed scFlow, an open-source analysis pipeline comprising i) the scFlow toolkit built in R with high levels of abstraction on top of popular single-cell analysis tools (e.g. Seurat, Monocle, Scater) and ii) nf-core/scFlow, a version-controlled, citable, NextFlow pipeline for the efficient orchestration of reproducible scRNA-seq analyses with scFlow (Di et al., 2017). Comprehensive reports with publication-quality figures detailing QC metrics, clustering, differential expression and pathway analyses are automatically generated. The modular nature of the scFlow toolkit provides the flexibility to specify alternate algorithms for key analytical steps while capturing analysis parameters comprehensively and generating interactive reports and publication-quality outputs. The nf-core/scFlow NextFlow workflow, which is engineered to follow strict best-practices guidelines of the nf-core community framework, enables “one-click” scRNA-seq analyses for users that apply easily specifiable analytical parameters and experimental design specifications to orchestrate reproducible and portable (computational infrastructure-independent) analyses inside containerized environments (Ewels et al., 2020). The extensibility and modular design of scFlow should enable future updates to incorporate new methods in the field. Below, we briefly summarize the core features of scFlow and its application to published single-cell datasets.

## Implementation

### Overview

scFlow is comprised of two components: i) an independent R package, scFlow, containing a toolkit for analysis of single-cell RNA sequencing data and ii) a Nextflow pipeline, nf-core/scflow, for orchestrating end-to-end, reproducible, automated and scalable single-cell analyses using the scFlow R package (Fig. 1).

**Figure 1:**
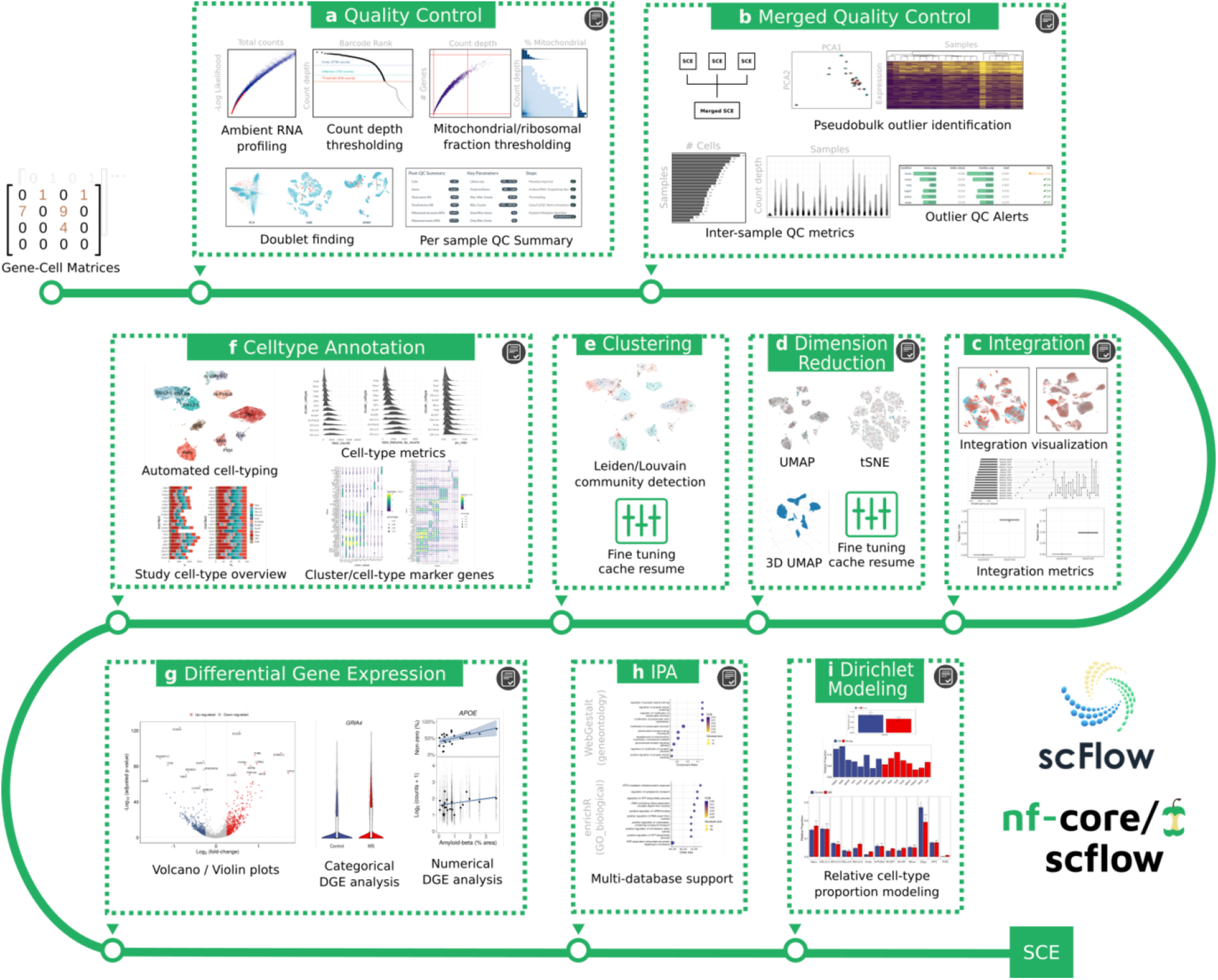
Single-cell analysis pipeline with nf-core/scflow using the scFlow toolkit. Gene-cell matrices from multi-sample case/control studies are analysed reproducibly across major analytical steps: (a) individual sample quality control including ambient RNA profiling, thresholding, and doublet/multiplet identification, (b) merged quality control including inter-sample quality metrics and sample outlier identification, (c) dataset integration with visualization and quantitative metrics of integration performance, (d) flexible dimension reduction with UMAP and/or tSNE, (e) clustering using Leiden/Louvain community detection, (f) automated cell-type annotation with rich cell-type metrics and marker gene characterization, (g) flexible differential gene expression for categorical and numerical dependent variables, (h) impacted pathway analysis with multiple methods and databases, and (i) Dirichlet modeling of cell-type composition changes. A high-quality, fully annotated, quality-controlled SingleCell-Experiment (SCE) object is output for additional downstream tertiary analyses. Interactive HTML reports are generated for each analytical step indicated (grey icon). Analyses are efficiently parallelized where relevant (steps a,g,h, and i) and all steps benefit from NextFlow cache enabling parameter tuning with pipeline resume particularly useful for dimension reduction (d) and clustering (e).

The scFlow R package is built to enable standardized workflows following best practices on top of popular single-cell R packages, including Seurat, Monocle, scater, emptyDrops, DoubletFinder, LIGER, and MAST (Hao et al., 2021; Cao et al., 2019; McCarthy et al., 2017; Lun et al., 2019; McGinnis et al., 2019; Welch et al., 2019). scFlow provides the ability to undertake common analytical tasks required by users that involve multiple tools with a single command (i.e. a higher level of abstraction). The Bioconductor SingleCellExperiment class (Amezquita et al., 2020) is utilized throughout, with the interconversion between package-specific object classes handled “under-the-hood” to perform analytical steps and return their results seamlessly. Analytical parameters are recorded comprehensively and made readily available to enable reproducible optimizations of analyses. Interactive HTML reports are generated for each stage of the analysis that describes algorithm performance metrics and provide publication-quality plots of a wide range of outputs, along with bibliographic citations for the analytical packages used. These reports thus provide the user with informative summaries of their specific analytical steps in ways that can highlight the impact of parameter choices and guide their revision when needed. The use of modular functions which receive and return a SingleCellExperiment object with relevant metadata appended allows new algorithms to be readily implemented. The following example illustrates a complete sample quality-control with default parameters using scFlow in R, including ambient RNA profiling, gene/cell annotation, thresholding, doublet/multiplet removal, and generation of an interactive HTML report with key plots: -

~~~
sce <- read_sparse_matrix(matrix_path) %>%
generate_sce(metadata) %>%
find_cells() %>%
annotate_sce() %>%
filter_sce() %>%
find_singlets() %>%
filter_sce() %>%
report_qc_sce()
~~~

### Analytical steps with scFlow

#### Quality-control

Initial quality-control is performed individually for each sample (Fig. 1a), using the post-demultiplexed sparse gene-cell counts matrix as input. Each sparse matrix is combined with unique sample metadata to generate the initial SingleCellExperiment (SCE) object. Ambient RNA profiling is performed optionally, using the EmptyDrops algorithm to flag and subsequently filter cellular barcodes which do not deviate from an ambient RNA expression profile representing cell-free transcripts (Lun et al., 2019). The SCE is subsequently annotated with rich gene and cell-level metrics and appended with key plots to guide parameter selection according to best practices, including barcode rank plots, and histograms of total counts, total features, and relative mitochondrial and ribosomal gene counts (Luecken and Theis, 2019). The ability to adaptively threshold cell metrics based on median absolute deviations enables consistent thresholding criteria to be applied across samples with different characteristics (e.g. between batches, across data from different species) to support integrative analyses.

The pipeline provides an option for submitting filtered post-QC SCE for doublet/multiplet detection using the DoubletFinder algorithm (McGinnis et al., 2019). Cells are embedded in reduced dimensional space using PCA, tSNE, and UMAP to facilitate visualization of putative non-singlets (which typically form isolated clusters or are embedded at the peripheries of major clusters) identified by the algorithm. A post-QC summary report brings relevant plots and algorithm performance metrics together to facilitate the joint consideration of QC covariates in univariate thresholding decisions, consistent with best practices (Luecken and Theis, 2019) (File 1).

#### Hosted file

`File_1_Zhou_et_al_human_dimis_qc_report.html` available at https://authorea.com/users/226952/articles/480342-scflow-a-scalable-and-reproducible-analysis-pipeline-for-single-cell-rna-sequencing-data

Following the merging of multiple post-QC samples, an additional post-merge QC step is applied to evaluate comparative metrics of sample quality (Fig. 1b). Firstly, a “bulk” RNA seq PCA plot of samples is generated by pseudobulking counts by sample, with an additional hierarchical clustering plot of binarized gene expressivity to highlight samples with a divergent feature space. Next, the total number of cells contributed by each sample is determined, and violin plots and interactive tables are generated for each user-specified cellular variable of interest (e.g. total counts, total features, relative mitochondrial counts, etc.), optionally stratified by experimental variables (e.g. batch). The tables additionally provide outlier warnings ([?]2σ) and alerts ([?]3σ) for each sample QC metric. Together, these results are collated in a post-merge QC report (File 2) both to guide the identification of putative sample-level outliers and any required revisions of QC parameters.

#### Hosted file

`File_2_Mathys_et_al_merged_report.html` available at https://authorea.com/users/226952/articles/480342-scflow-a-scalable-and-reproducible-analysis-pipeline-for-single-cell-rna-sequencing-data

#### Integration and dimensionality reduction

Latent metagene factors representing shared features of cell identity across different experimental samples can be generated using the linked inference of genomic experimental relationships (LIGER) algorithm (Fig. 1c) (Welch et al., 2019). Providing these latent factors as inputs in place of principal components for dimensionality reduction can improve dataset integration. Dimensionality reduction then is performed using the uniform manifold approximation and projection (UMAP) or t-distributed stochastic neighbour embedding (tSNE) algorithms to generate 2D or 3D embeddings (Fig. 1) (Kobak and Berens, 2019; Becht et al., 2018). The performance of any dataset integrations can subsequently be assessed across user-specified experimental covariates (e.g. batch, sex, disease) using a combination of juxtaposed reduced dimension plots with and without integration and quantitative scores of cell mixing using ‘rejection rates’ from the k-nearest-neighbor batch-effect (kBET) algorithm (üttner2019**?**). These results, together with details of the latent factors generated by LIGER (e.g. UpSet plots of dataset participation), are brought together in an integration report that serves to characterise performance of the integration algorithm and thus can be used to guide revisions of integration and dimensionality reduction parameters (Fig. 1c) (File 3).

#### Hosted file

`File_3_Ximerakis_et_al_integrate_report.html` available at https://authorea.com/users/226952/articles/480342-scflow-a-scalable-and-reproducible-analysi-pipeline-for-single-cell-rna-sequencing-data

#### Clustering and cell-type annotation

Cell clusters are identified with the Leiden or Louvain community detection algorithms implemented in Monocle using the UMAP or tSNE embeddings as inputs (Fig. 1e) (Traag et al., 2019; Trapnell et al., 2014). Following clustering, automated cell-type prediction is performed on cell clusters using the expression weighted cell type enrichment (EWCE) algorithm against reference datasets previously generated with EWCE (Fig. 1f) (Skene and Grant, 2016). Detailed cell-type metrics are subsequently generated, including plots of the relative proportions of cell-types by user-specified experimental variables (e.g. sample, diagnosis), histograms of user-specified cell metrics (e.g. total counts, total features, relative mitochondrial counts) and detailed dot-plots and interactive tables of cluster and cell-type marker genes generated using Monocle (Trapnell et al., 2014). These results are collated into a comprehensive cell-type metrics report (File 4), enabling multi-parametric characterisation of cell types and guiding any subsequent manual revisions of cell-type labels or clustering parameters that may be demanded.

#### Hosted file

`File_4_Mathys_et_al_celltype_metrics_report.html` available at https://authorea.com/users/226952/articles/480342-scflow-a-scalable-and-reproducible-analysis-pipeline-for-single-cell-rna-sequencing-data

#### Differential gene expression and impacted pathway analysis

Differential gene expression (DGE) within cell-types can be evaluated for both categorical (e.g. diagnosis) and numerical (e.g. age, pathology scores) dependent variables while accommodating complex experimental designs and controlling for covariates (Fig. 1g). A pre-processing step enables optional filtering of genes based on expressivity, pseudobulking, input matrix transformation (e.g. Log2, CPM), and co-variate scaling and centering. The default DGE method in scFlow is a generalized linear mixed model (GLMM) with a random effect (RE) term (e.g., to account for correlations within individual samples) as implemented within the model-based analysis of single-cell transcriptomics (MAST) algorithm (Zimmerman et al., 2021; Finak et al., 2015). An interactive DGE HTML report with a volcano plot and searchable tables is generated, including details of model parameters, inputs, and outputs (File 5).

#### Hosted file

`File_5_Mathys_et_al_Oligo_MASTZLM_Control_vs_pathological_diagnosisAD_de_report.html` available at https://authorea.com/users/226952/articles/480342-scflow-a-scalable-and-reproducible-analysis-pipeline-for-single-cell-rna-sequencing-data

Impacted pathway analysis (IPA) is performed on DGE tables to identify enrichment of differentially expressed genes in specific pathways (Fig. 1h). Comprehensive methods and databases available within the enrichR (Kuleshov et al., 2016), ROntoTools (Khatri et al., 2007), and WebGestaltR (Liao et al., 2019) packages can be used simultaneously for the generation of an interactive HTML report including dot-plots for the top enriched pathways and searchable tables of results across different methods (File 6).

#### Hosted file

`File_6_Mathys_et_al_Oligo_MASTZLM_Control_vs_pathological_diagnosisAD_DE_ipa_report.html` available at https://authorea.com/users/226952/articles/480342-scflow-a-scalable-and-reproducible-analysis-pipeline-for-single-cell-rna-sequencing-data

#### Modeling of relative cell-type proportions

Statistically significant changes in cell-type composition across categorical dependent variables (e.g. case vs control) can be examined using a Dirichlet-multinomial regression model, which accounts for dependencies in cell-type proportions within samples (Fig. 1i) (Smillie et al., 2019). Adjusted p-values and plots of relative abundance are generated for each cell-type and collated in an HTML report together with composition matrices used in model generation (File 7).

#### Hosted file

`File_7_Ximerakis_et_al_dirichlet_report.html` available at https://authorea.com/users/226952/articles/480342-scflow-a-scalable-and-reproducible-analysis-pipeline-for-single-cell-rna-sequencing-data

### Pipeline orchestration with nf-core/scflow

#### Overview

We built the nf-core/scflow pipeline using Nextflow within the nf-core framework to enable standardized, portable, and reproducible analyses of case/control single-cell RNA sequencing data (Ewels et al., 2020). Pipelines built using Nextflow inherit its portability, native support for container technologies, and features including cache-based pipeline resume capability and amenability to live-monitoring (Di et al., 2017). The nf-core framework provides a means to produce high-quality, best-practices analysis pipelines with Nextflow which are ready for deployment across all institutions and research facilities (Ewels et al., 2020).

#### Workflow

The codebase for both the scFlow R package toolkit and the nf-core/scflow pipeline are stored in open-source GitHub repositories (Fig. 2). Both repositories are version controlled and utilize continuous integration (CI) workflows to ensure code updates pass build and functionality tests. In addition, updates to nf-core/scflow trigger an automated CI action to validate that the analysis of a small case/control dataset runs to completion without errors. Version updates to the scFlow R package trigger a CI action to build a new version-tagged Docker image which is uploaded to a Docker registry. This image is built from a Dockerfile specification which additionally installs the complete set of software dependencies, including 414 versioned R packages and additional system-level dependencies (scFlow 0.7.1, see supplemental data).

**Figure 2:**
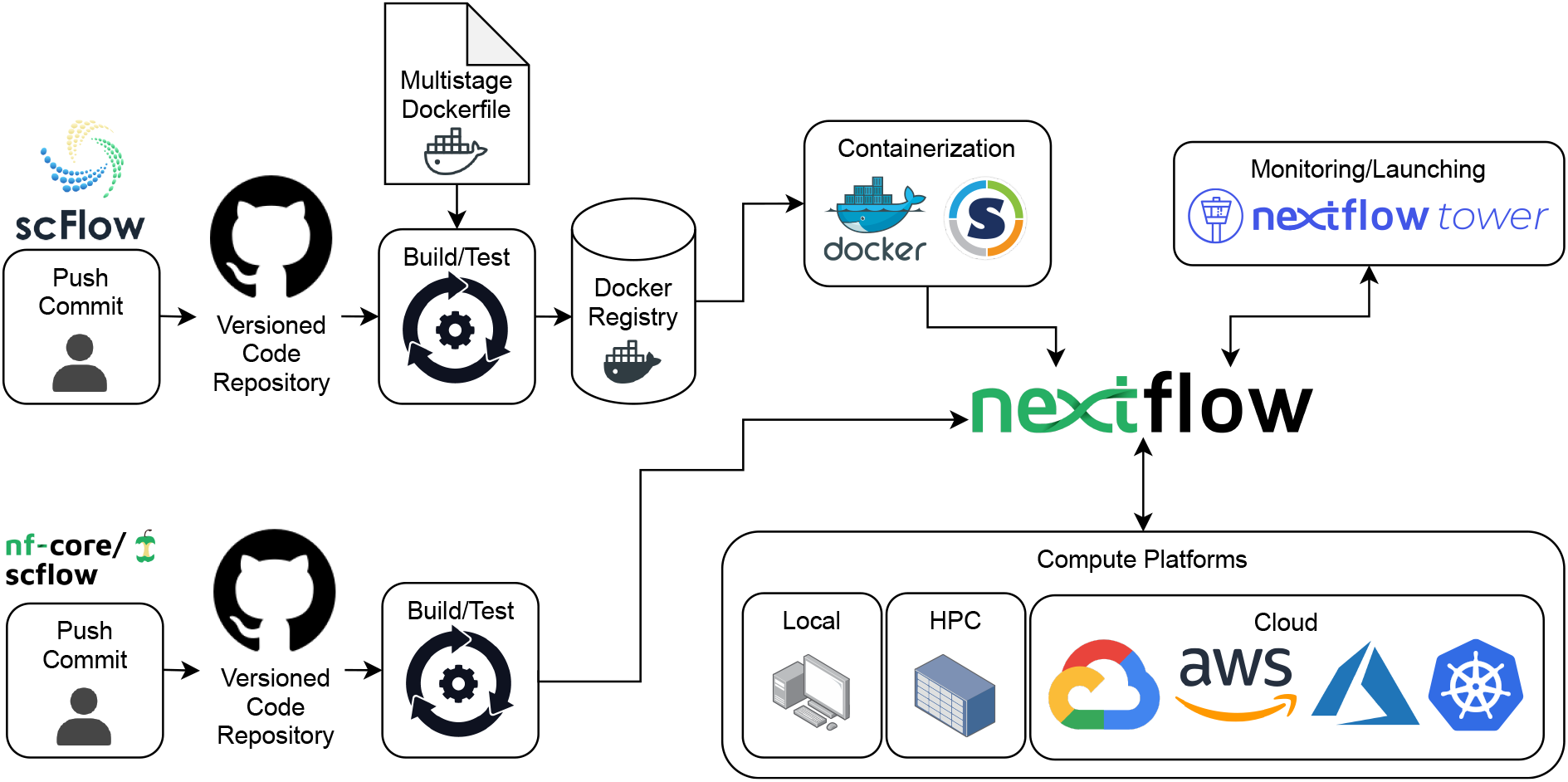
Workflow for scFlow and nf-core/scflow. Open-source code for both scFlow and nf-core/scflow is version-controlled and stored in GitHub repositories with continuous integration (CI) to build and test updated code. Container images including software dependencies are automatically built on version updates and uploaded to a Docker registry. Pipeline runs with nf-core/scflow utilize containerized environments using Docker/Singularity to perform analyses reproducibly across diverse compute infrastructure including local workstations, high- performance clusters (HPC), or Cloud services including Google Cloud, Amazon Web Services, Microsoft Azure, and Kubernetes. Real-time monitoring and optional launching of pipeline runs can be performed using NextFlow Tower.

The execution of an nf-core/scFlow pipeline run automatically retrieves the correct version of the Docker image from the Docker registry and generates reproducible containerized analysis environments for each analytical process using Docker or Singularity. Analyses are performed on the compute platform preferred by the user given the potential for implementation on local, high-performance computing cluster (HPC) or in Cloud based environments (Di et al., 2017) (Fig. 2). Live-monitoring of pipeline progress is possible using Nextflow Tower [https://tower.nf/], a hosted and open-source solution providing live statistics on resource usage (e.g. CPU, RAM, IO, time) and cost (for Cloud analyses). Pipeline runs can also be optionally launched directly from within the Nextflow Tower GUI.

#### Executing a nf-core/scflow pipeline run

A pipeline run with nf-core/scflow requires three inputs: (1) a two-column manifest file with paths to gene-cell matrices and a unique sample key; (2) a sample sheet with sample information for each input matrix in the manifest file; and, (3) a parameters configuration file (documentation for each parameter is available at https://nf-co.re/scflow/dev/parameters). A complete, automated, scalable, and reproducible case-control analysis following the steps in Figure 1 can then be performed with a single line of code: -

~~~
nextflow run nf-core/scflow \
--manifest Manifest.tsv \
--input Samplesheet.tsv \
-c scflow_params.config \
-profile local
~~~

Switching from a local workstation analysis to a Cloud based analysis can be achieved simply by changing the profile parameter. For example, a Google Cloud analysis with automated staging of input matrices from Cloud storage (e.g. a Google Storage Bucket) can be achieved using `-profile gcp`. Additionally, pre-configured institutional profiles for a range of university and research institution HPC systems are readily available via nf-core [https://github.com/nf-core/configs].

During an nf-core/scflow run, comprehensive pipeline outputs are generated including flat-file tables, images, and interactive HTML reports. As Nextflow utilizes an intelligent cache based on hashed inputs to each analytical task, the pipeline can be stopped at any time, parameters adjusted, and the pipeline resumed with the addition of the ‘`-resume`’ option. As only tasks downstream of the changed parameters are affected and re-run, parameter optimization is both simplified and accelerated, particularly for the early steps of dimensionality reduction, clustering, and optional revision of automated cell-type annotations.

## METHODS

### Ambient RNA profiling

Our implementation of EmptyDrops includes the default options with the emptyDrops R package, with the following additions. The threshold of UMI counts above which a cellular barcode will be retained can optionally be determined based on a quantile approach as described previously and implemented in the CellRanger software by 10X Genomics (Zheng et al., 2017). This ‘auto’ option for the retain parameter retains all barcodes with *>*10% of the counts in the top nth barcodes, where n is 1% of the expected recovered cell count specified by the ‘expect - cells’ parameter. The distribution of p-values for presumed ambient barcodes is evaluated for uniformity – as expected under the null-hypothesis – using a Kolmogorov-Smirnov test. Default emptyDrops parameters used by scFlow are: lower=100, retain=‘auto’, expect cells=3000, and niters=30000.

### Thresholding

For thresholds determined adaptively, a user-specified number of median absolute deviations (nMADs) is applied using the Scater package (McCarthy et al., 2017) as previously described (Lun et al., 2016b):

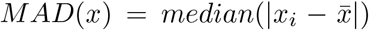. Default thresholding parameters used by scFlow are: min_library_size=100, max_library_size=‘adaptive’, min_features=100, max_features=‘adaptive’, max_mito=‘adaptive’, min_ribo=0, max_ribo=1, min_counts=2, min_cells=2, drop_unmapped=TRUE, drop_mito=TRUE, drop_ribo=FALSE, and nmads=4.0. Outliers for inter-sample post-merge quality-control metrics are determined based on standard deviation (σ) across samples with warnings provided at the [?]2σ level and alerts at the [?]3σ level.

### Pseudobulking

Pseudobulking is performed by summation of counts by sample as previously described (Lun et al., 2016a). For computational efficiency, the calculations are performed using matrix multiplication where rows (gene counts) are multiplied by columns of a sample annotation model matrix: *c*_*ik*_ = *a*_*ij*_*b*_*jk*_.

### Doublet/multiplet detection

The DoubletFinder algorithm is implemented essentially as described using the DoubletFinder R package (McGinnis et al., 2019) with the following additions. A fixed doublet rate, or alternatively, a doublets-per-thousand-cells increment (‘dpk’ parameter) can be set to scale the doublet rate with the number of cells considered, as recommended by 10X Genomics. The ‘pK’ parameter can be fixed or determined following a parameter sweep to identify the ‘BCmetric’ maxima across a range of ‘pK’ values, as described in the DoubletFinder vignette. Default parameters used for DoubletFinder in scFlow are: pca_dims=20, var_features=2000, dpk=8, and pK=0.02.

### Dataset integration

LIGER (‘rliger’ package) was used for dataset integration which uses an integrative non-negative matrix factorization (iNMF) method to identify shared and dataset-specific factors. The latent metagene factors are generated as previously described (Welch 2019, Liu 2020). Four preprocessing steps are applied: (1) normalization for UMIs per cell using ‘rliger::normalize’, (2) subsetting the most variable genes for each dataset using ‘rliger::selectGenes’, (3) scaling by root-mean-square across cells using ‘rliger::scaleNotCenter’ to ensure different genes have the same variance, and (4) filtering of non-expressive genes. For integration, we use the union of the top ‘num genes’ variable genes from each dataset. To ensure that the union is not significantly skewed towards a specific dataset(s), we identify possible outlying dataset(s) using Venn and UpSet diagrams generated by ‘nVennR::plotVenn’ (Perez-Silva 2018) and ‘UpSetR::upset’ (Conway 2017), respectively. The shared and dataset-specific factors are subsequently generated from the normalized and scaled inputs using iNMF with the ‘rliger::optimizeALS’ function. Finally, the ‘rliger::quantile norm’ function is applied to integrate the datasets together using a maximum-factor assignment followed by refinement using a k-nearest neighbours (KNN) graph. Default parameters used for LIGER are: take_gene_union=FALSE, remove_missing=TRUE, num_genes=3000, combine=“union”, capitalize=FALSE, use_cols=TRUE, k=30, lambda=5.0, thresh=0.0001, max_iters=100, nrep=1, rand_seed=1, knn_k=20, ref_dataset=NULL, min_-cells=2, quantiles=50, resolution=1 and centre=FALSE. Performance of the integration algorithm is evaluated quantitatively using the kBET algorithm essentially as previously described (Büttner 2019). A low ‘rejection rate’ determined by kBET indicates cells from different batches (and/or other user-defined categorical covariates) are well-mixed.

### Dimensionality reduction

The top *m* principal components are calculated based on highly variable genes using ‘Seurat::RunPCA’ for Seurat based sub-workflows, otherwise ‘monocle3::preprocess cds’ is used. Embeddings for tSNE are generated using ‘Seurat::RunTSNE’ for Seurat based sub-workflows, otherwise, Jesse Krijthe’s ‘Rtsne::Rtsne’ implementation of Van der Maaten’s Barnes-Hut algorithm is used [https://github.com/jkrijthe/Rtsne]. Default parameters for tS-NE are: dims=2, initial_dims=30, perplexity=50, theta=0.5, stop_lying_iter=250, mom_-switch_iter=250, max_iter= 1000, pca_center=TRUE, pca_scale=FALSE, normalize=TRUE, momentum=0.5, final_momentum=0.8, eta=1000, and exaggeration_factor=12. Embeddings for UMAP are generated using ‘Seurat::RunUMAP’ for Seurat based sub-workflows, otherwise, James Melville’s ‘uwot::umap’ implementation of the UMAP algorithm is used [https://github.com/jlmelville/uwot]. Default parameters used for UMAP are: pca_dims=30, n_neighbors=35, n_components=2, init= ‘spectral’, metric=‘euclidean’, n_epochs=200, learning_rate=1, min_dist=0.4, spread=0.85, set_op_mix_ratio=1, local_connectivity=1, repulsion_-strength=1, negative_sample_rate=5, and fast_sgd=FALSE.

### Clustering

Clustering of cells using the Louvain or Leiden community detection algorithms is performed using the ‘monocle3::cluster_cells’ function, with the modified ability to cluster on any named reducedDims matrix of the SingleCellExperiment object (e.g. UMAP embeddings from LIGER generated latent factors, UMAP_Liger). Default parameters are set to cluster_method=‘leiden’, res=1e-5, k=100, and Louvain_iter=1.

### Cell-type annotation

Automated cell-type annotation is performed using the expression weighted cell type enrichment (EWCE) package essentially as previously described (Skene and Grant, 2016). Reference datasets containing annotated cell-types are first processed using EWCE to produce cell-type data (‘CTD’) files comprised of cell-type-specific transcriptional signatures. In our analyses, we have used ‘CTD’ files generated from the Allen human brain atlas (Hodge et al., 2019) and a mouse brain dataset (Zeisel et al., 2015). The top 10% most specific genes are used as marker genes for each cell-type. Up to *m* (default: 10000) cells sampled from the numbered Louvain/Leiden clusters are evaluated for statistical enrichment in target gene lists of length *n* from the reference ‘CTD’ against a background probability distribution generated by 1000 permutations of random background gene lists of length *n*. Each cluster is subsequently annotated with the highest scoring (lowest adjusted p-value) cell-type and the complete set of results are returned with the SingleCellExperiment metadata.

### Differential gene expression

For DGE, a pre-processing step is first performed to subset genes based on expressivity within a specific cell-type (default: [?]1 count in [?]10% of cells). Next, the percentage of variance in gene expression explained by inter-sample variation within a reference class (e.g. healthy/control) can optionally be calculated using the ‘scater::getVarianceExplained’ function (McCarthy et al., 2017). These values are ranked and appended to the output DGE table as an additional sense check and optional gene list filtering criterion. The proportion of genes detected in each cell is then calculated and scaled to obtain the cellular detection rate (CDR), as previously described (Finak et al., 2015):

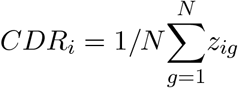

Numerical predictors (e.g. age, quantitative histopathology measure) can be scaled and centered prior to model fitting. Optionally, pseudobulking also can be performed, as described above (). After pre-processing, DGE models with MAST are performed essentially as previously described (Finak et al., 2015). A log2(TPM + 1) expression matrix is calculated from the raw counts matrix, and a two-part (i.e., including a discrete logistic regression component for expression rate and a continuous Gaussian component conditioned on each cell expressing a gene) generalized regression model is fit independently for each gene. The CDR is included as a covariate alongside additional user-specified experimental covariates, which can include, for example, the individual sample as a random effect (Zimmerman et al., 2021). False-discovery rate (FDR) adjusted p-values are determined using the Benjamini & Hochberg method.

### Impacted pathway analysis

Enrichment of gene lists in pathways are evaluated using methods encompassing Over Representation Analysis (ORA) (Khatri et al., 2012), Gene Set Enrichment Analysis (GSEA) (Subramanian et al., 2005), and Network Topology-based Analysis (Wang et al., 2017). Databases and methods from one or more of the R packages WebGestaltR (Liao et al., 2019), ROntoTools (Mitrea et al., 2013), and enrichR (Chen et al., 2013) are applied as previously described, and can be queried simultaneously. Results are returned as standard tool output tables and dot plots of enrichment/odds ratio vs adjusted p-values (FDR) for the top *n* pathways are generated and collated into an IPA HTML report.

### Dirichlet modeling of cell-type composition

To identify statistical differences in cell-type proportions between categorical dependent variables (e.g. case vs control), a Dirichlet multinomial regression is performed (Smillie et al., 2019). A sample (rows) by cell-types (columns) matrix of cell numbers is generated and normalized to relative proportions (0, 1) such that the sum of proportions of each cell-type *c* in sample *y* equals one: 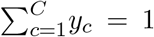. If extreme values of 0 or 1 are present, a transformation is applied using the DirichletReg R package to shrink values away from these extremes by transforming each component y of Y by computing *y*\* = [*y*(*n −* 1) + 1/*d*]/*n* where n is the number of observations in Y, as implemented in the ‘DR_data’ function in the R package DirichletReg (Maier, 2014). The “common” model (counts ∼ dependent_variable) is fit using the ‘DirichletReg::DirichReg’ function and p-values are extracted. Bar plots for each cell-type are generated and collated with input and output tables for the cell-type proportions HTML report.

### Dataset pre-processing

Inputs for scFlow are standardized sparse-matrices generated by widely-used pipelines (e.g. Cell Ranger) for processing, reference genome mapping, and de-multiplexing of raw single-cell sequencing data (Zheng et al., 2017). As public datasets vary with respect to data deposition format, custom scripts were required for each of the four analysed datasets to (a) pre-process matrices into standard per-sample gene-cell counts matrices and/or (b) build a sample sheet with pertinent experimental data attached. Raw gene-cell count matrix and sample-level metadata for (Mathys et al., 2019) and the human dataset for (Zhou et al., 2020) were downloaded from the AD Knowledge Portal [https://www.synapse.org] (Synapse ID: syn18485175) and the count matrix for the respective dataset was split into per sample gene-cell count matrices. Mouse datasets (Zhou et al., 2020)(Ximerakis et al., 2019) were downloaded from GEO (https://www.ncbi.nlm.nih.gov/geo/) (ID:GSE140511 and GSE129788, respectively) and split into per sample gene-cell count matrices. The feature names for gene-cell count matrices from (Ximerakis et al., 2019) were mouse gene symbols which were first converted to mouse Ensembl IDs. All data preprocessing scripts are available at https://github.com/combiz/scFlow_Supplementary.

### Nextflow

The nf-core/scflow pipeline was coded in Nextflow with domain-specific language 2 (DSL2) according to nf-core guidelines. The major analytical steps outlined in Figure 1 are performed across Nextflow processes implemented in DSL2 modules as detailed in (S**??**). Included are details of the underlying scFlow functions utilized for each process, an overview of process outputs, and parallelization support (i.e. simultaneous analysis of multiple samples/models across multiple independent compute instances/jobs). These processes represent modular units of pipeline execution in Nextflow, simplifying the modification of individual pipeline steps, and allowing process-level resource allocation. Detailed information on pipeline usage, parameters, and outputs are provided in the nf-core/scflow documentation online [https://nf-co.re/scflow].

### Software availability

The code for the scFlow R package is available in a GitHub repository [https://github.com/combiz/scflow] with associated function documentation at https://combiz.github.io/scFlow. The code for the nf-core/scflow pipeline is available in a GitHub repository [https://github.com/nf-core/scflow] with pipeline documentation at https://nf-co.re/scflow. A general usage manual is available at https://combiz.github.io/scflow-manual/. All code is open-source and available under the GNU General Public License v3.0 (GPL-3).

## Results

To demonstrate the performance and flexibility of scFlow for automated case-control sc/snRNA seq analyses, four previously published datasets were retrieved from online repositories, pre-processed, and submitted to nf-core/scflow for analysis with a single line of code (Mathys et al., 2019; Ximerakis et al., 2019; Zhou et al., 2020). These studies encompass samples from both human and mouse species, include both single-cell and single-nuclei data, span a range of samples per study (12 - 48), and each represent a different type of experimental design with different confounds and variables of interest (Table 1).

**Table 1:**
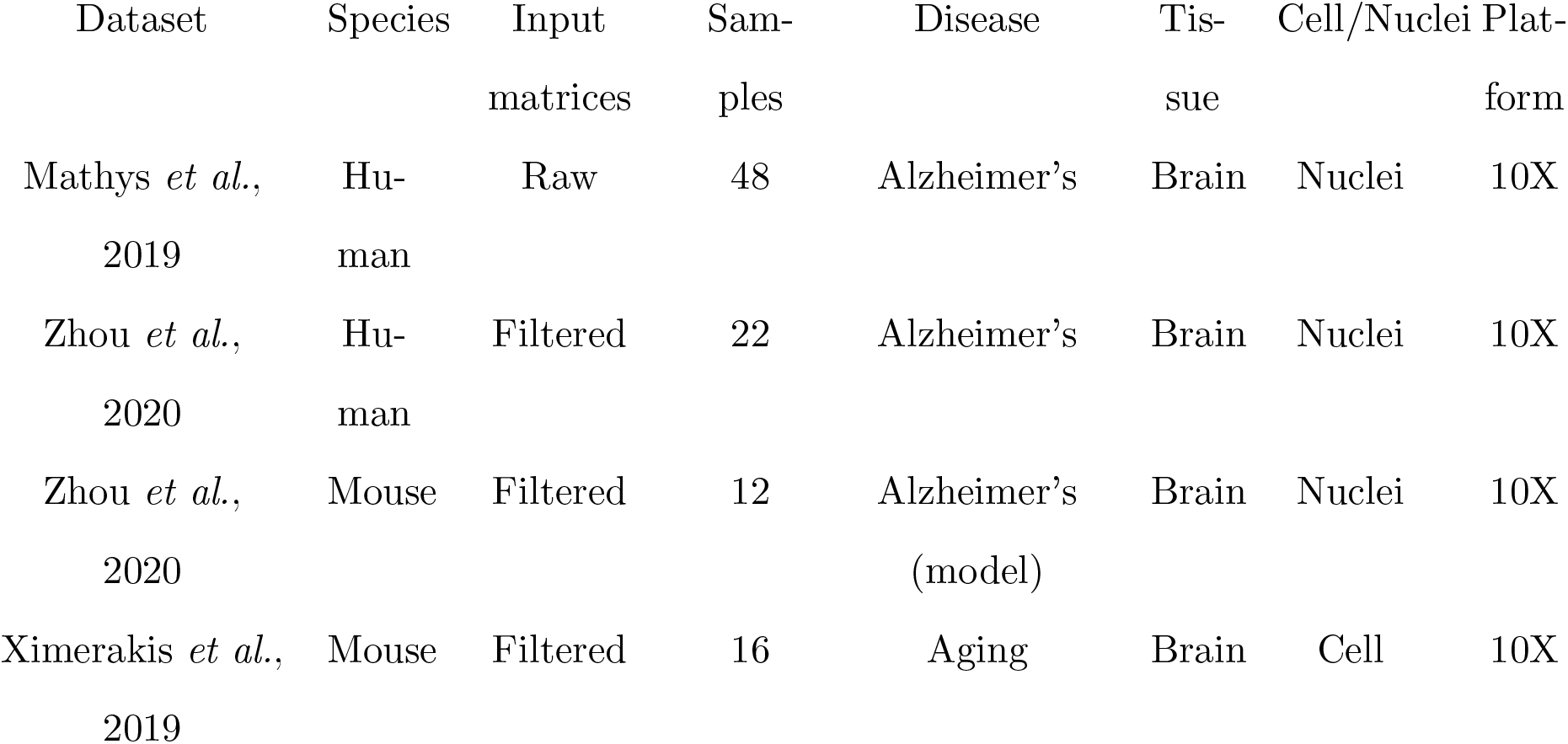
Characteristics of individual datasets analysed in this study with nf-core/scflow.

Selected cell-level quality-control metrics and cell/gene-level inclusion and exclusion QC checks presented here highlight the valuable quality-control data captured by the pipeline (Figure 3). The complete set of pipeline QC outputs are included in the supplemental materials (e.g. per-sample QC reports, study post-merge reports, QC metrics summary table). The extensive variation between samples within - and across - studies that is apparent illustrates the importance of tailored thresholding (e.g. minimum and maximum counts, features, relative mitochondrial counts, etc.) and the potential benefits of identifying sample-level outliers.

**Figure 3:**
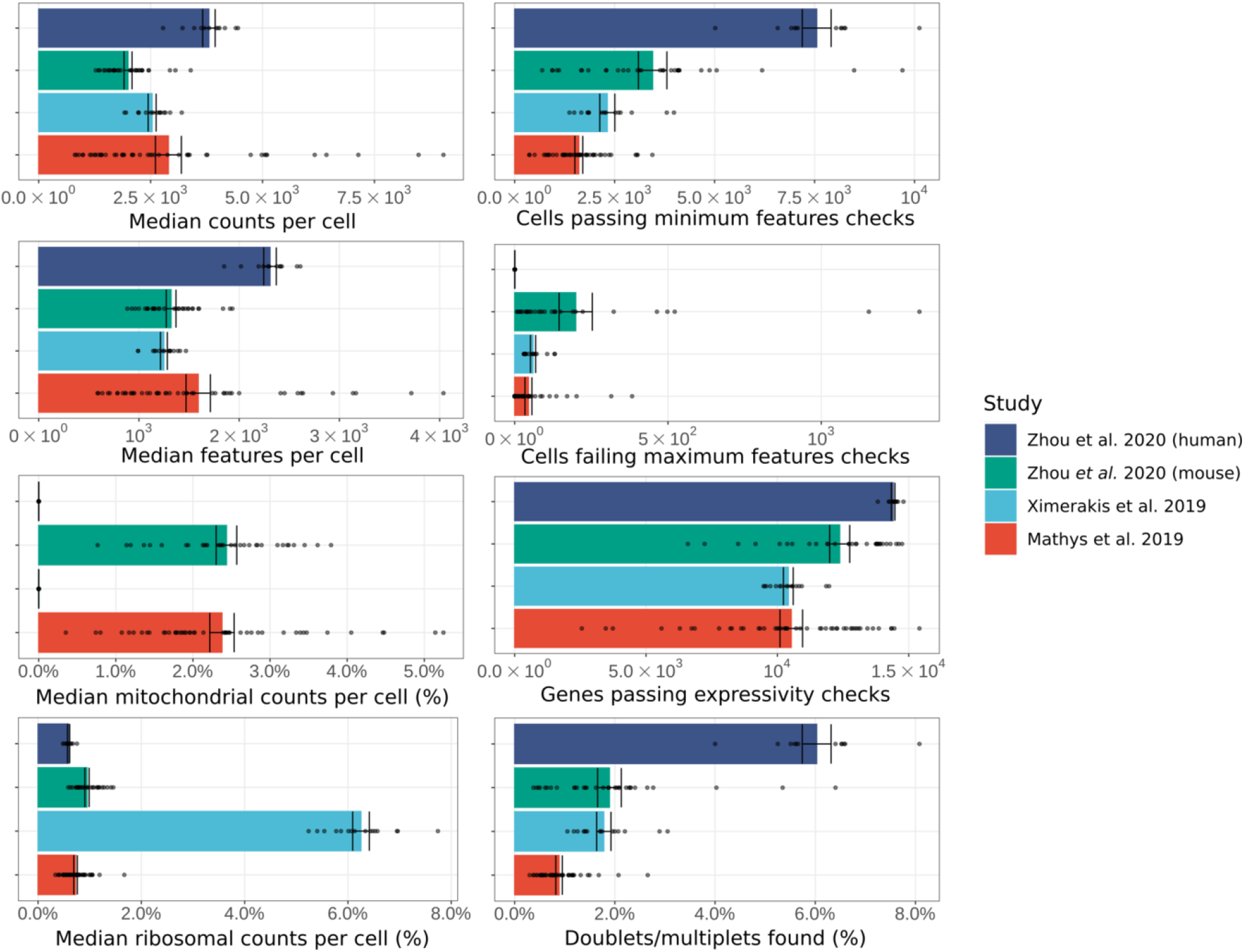
Selected quality-control metrics in 98 samples across four datasets. The mean (bar ±SEM) of the sample-level medians (points) of four key cell-level metrics – total counts, total genes, relative mitochondrial counts, and relative ribosomal counts – are presented for each of the four analysed datasets (colours) in the left column. The right column includes examples of cell- and gene-level quality control inclusion and exclusion checks used for filtering of input matrices for downstream analyses.

The UMAP embeddings with cell-type annotations show a good separation (global distance) between major cell-types (e.g. oligodendrocytes and astrocytes) with relative proximity of related cell-types (e.g. neuronal sub-types) for each of the four datasets, as expected (Figure 4a). Additionally, the cell-type markers identified by the pipeline are consistent with known markers for the cell-types (see supplemental data, cell-type metrics reports). The UMAP embeddings for all four datasets were generated from latent metagene factors computed by LIGER. This integration approach leads to UMAP embeddings that are less driven by known sources of variation in the data (e.g. diagnosis, age, genotype). This is demonstrated both visually – by contrast to a unintegrated (left) UMAP (Figure 4b) – and by a reduced kBET ‘rejection rate’, reflecting improved cell mixing (Figure 4c). Improved integration across multiple additional sources of sample-level variance (e.g. individual, sex) are also evident (see supplemental data, integration reports). Together these provide evidence that integration of the data was effective, with a greater contribution of shared, relative to sample-specific, factors to the separation of cells in reduced dimensional space.

**Figure 4:**
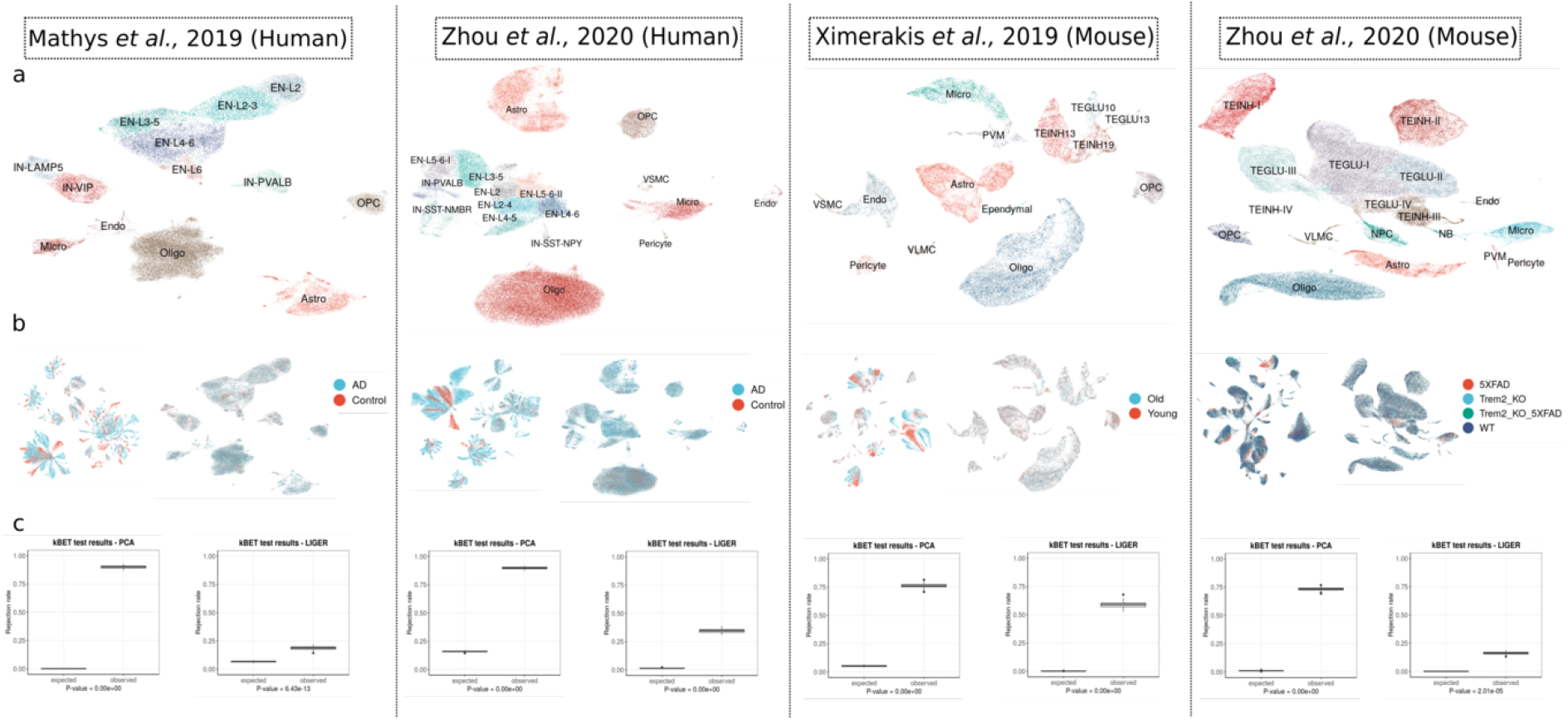
Cell-type annotation and key integration results from the analysis of four datasets with the nf-core/scflow pipeline. For each study, a) UMAP plot of the final clusters with their cell-type annotations, b) UMAP of an unintegrated (left) and LIGER-integrated (right) dataset highlighting the categorical variable of interest, c) box plots of expected and observed kBET ‘rejection rates’ from unintegrated (left) and LIGER-integrated (right) UMAPs.

The relative proportion of cell-types in each study, further stratified by the major dependent variable of interest, is summarized in Figure 5a. Differential gene expression was evaluated in each cell-type using a mixed-model in MAST with a random effect for individual. The number of differentially expressed genes identified as up-regulated and down-regulated for each cell-type are highlighted (Figure 4b). For a selected cell-type from each study, the number, significance (adjusted p-value), and magnitude (fold-change) of evaluated genes are illustrated in a volcano plot (Figure 4c). Although an in-depth contrast of our results with those in the original studies is beyond the scope of this manuscript, the identification of the canonical Alzheimer’s disease implicated gene ‘Apoe’ in the microglia of mouse cells in the Alzheimer’s mouse dataset from Zhou *et al*. provides an example of the potential for insight discovery using our pipeline. Overall, these results, associated with similarly identified cells and derived using the same, well-controlled analytical pipeline and parameters, highlight the clear differences between differentially expressed gene sets from different studies and tissue types. By doing so, they also allow more confident generalisations regarding those features that are reproducible (e.g., the greater complexity and numbers of significantly differentially expressed genes in the rapidly isolated single-cell mouse brain transcriptomes relative to those in the human single nuclear transcriptomes from post mortem brain tissue).

**Figure 5:**
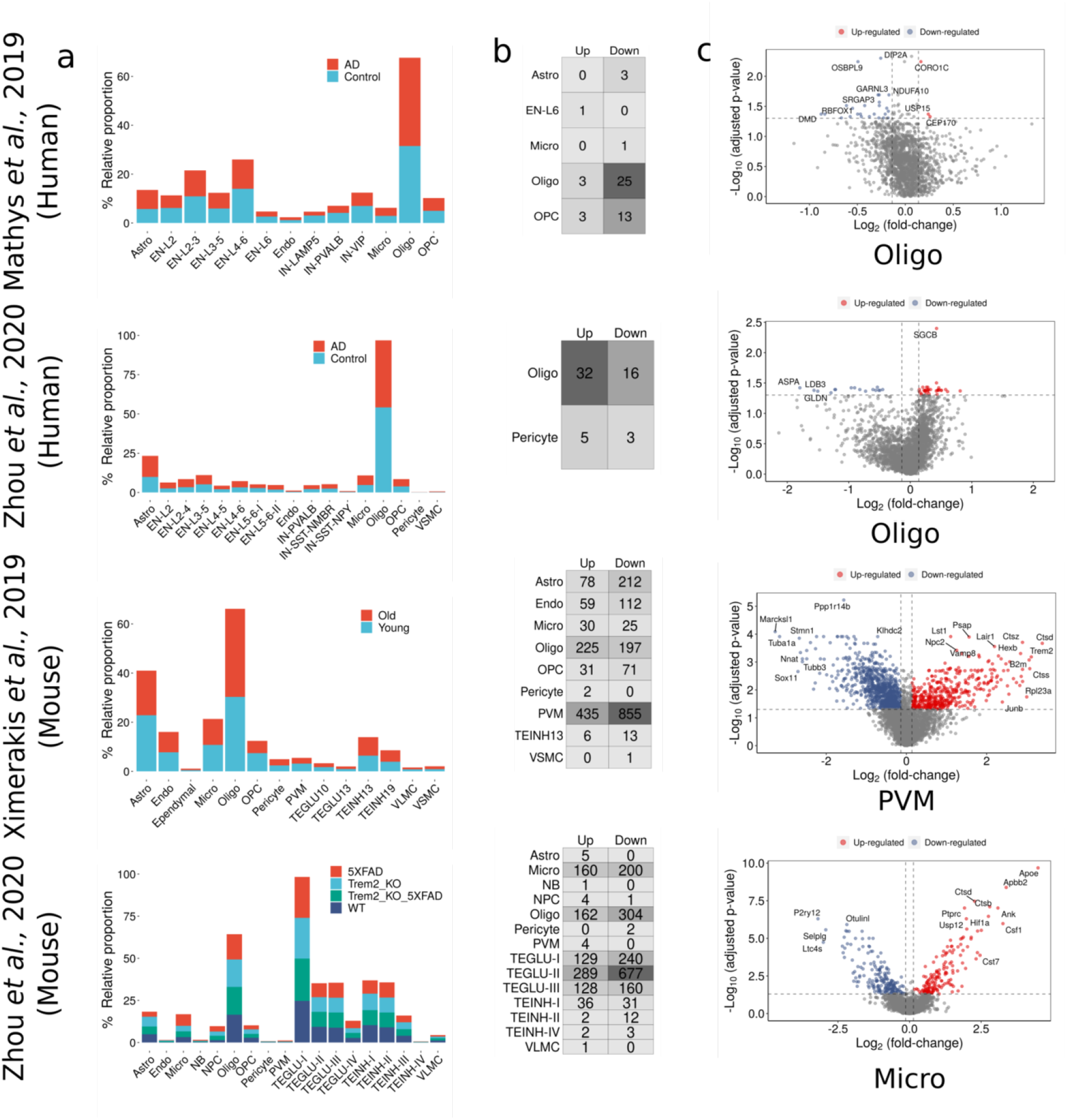
Relative cell-type proportions and summary of differential expression analysis results. a) Bar plots showing the relative proportions of cell-types between the categorical variables of interest for each of the datasets, b) Numbers of statistically significant up and down- regulated genes per cell-type; cell-types for which no differential expressed genes were found were omitted, c) Volcano plots of differentially expressed genes for the specified cell types of the four datasets.

## Discussion

The use of pipelines for the analysis of large datasets involving multiple, complex analytical steps is essential to achieve reproducible results. The wide range of alternative tools available for most analytical steps of single-cell RNA sequencing, combined with the different experimental questions and confounds particular to each dataset, often leads to project-specific code. These would typically require the manual revision of code to analyze a new dataset or to utilize an alternative algorithm for an analytical step. The scFlow toolkit and nf-core/scflow pipeline address this by implementing standardized, modular code: the flexibility to handle complex experimental designs and apply alternative algorithms are handled at the level of parameter specification. This modular approach also lends itself well to extensibility, as new tools in the field may be readily incorporated for an individual analytical task. The decoupling of analysis logic from resource allocation by Nextflow provides portability and scalability, with nf-core/scflow ready to run on local workstations, HPC environments, and Cloud services including Google Cloud, Amazon Web Services, and Microsoft Azure. This scalability will allow scFlow to keep pace with the burgeoning scale of single-cell RNA sequencing datasets.

The use of containerization technology by nf-core/scflow provides a consistent computing environment to ensure that the complex software and system dependencies used for analysis are comprehensively captured and are re-usable. Taken together with the version-control of pipeline code, and the generation of a citable unique digital object identifier (DOI) via nf-core for each versioned update to the pipeline, there is reassurance both of the reproducibility and the citability of an analysis.

We expect that the ease-of-use of nf-core/scflow and its flexibility to integrate datasets should be particularly useful for case-control and joint studies, including cell-atlas projects where data may be generated at different sites using different scRNA-seq protocols. The ability to adaptively threshold samples and evaluate inter-sample quality-control metrics can inform sample inclusion/exclusion criteria and potentially greatly improve the quality of such data resources.

In summary, the scFlow toolkit and nf-core/scflow pipeline provide a robust and easy-to-use analysis approach, leveraging the best scRNA-seq analysis tools in the R ecosystem with stateof-the-art data science to provide scalable, reproducible, and extensible analyses of scRNA-seq data.

## Supporting information

Supplementary Report Files 1 to 7

## Acknowledgements

CK is grateful to the Imperial College London NIHR Biomedical Research Centre Brain Sciences Theme for funding for his work. NS acknowledges support from UKRI Future Leaders Fellow-ship [grant number MR/T04327X/1]. PMM acknowledges generous personal support from the Edmond J Safra Foundation and Lily Safra and an NIHR Senior Investigator Award. This work was supported by the UK Dementia Research Institute, which receives its funding from UK DRI Ltd., funded by the UK Medical Research Council, Alzheimer’s Society and Alzheimer’s Research UK, and the Imperial College London NIHR Biomedical Research Centre.

We are grateful for helpful comments from members of the Nextflow and nf-core teams, in particular Paolo Di Tommaso, Philip A. Ewels, Harshil Patel, Alexander Peltzer, and Maxime Ulysse Garcia, and lab members including Johanna Jackson, Amy Smith, Karen Davey, and Stergios Tsartsalis.

## Declarations

PMM has received consultancy fees from Roche, Adelphi Communications, Celgene, Neurodiem and Medscape. He has received honoraria or speakers’ fees from Novartis and Biogen and has received research or educational funds from Biogen, Novartis and GlaxoSmithKline.

## Supplementary Material

### Hosted file

`Supplementary_Table_2_Process_Summary.xlsx` available at https://authorea.com/users/226952/articles/480342-scflow-a-scalable-and-reproducible-analysis-pipeline-for-single-cell-rna-sequencing-data

### Hosted file

`Supplemental_software_versions.tsv` available at https://authorea.com/users/226952/articles/480342-scflow-a-scalable-and-reproducible-analysis-pipeline-for-single-cell-rna-sequencing-data

### Hosted file

`Supplemental_Reports_Info.md` available at https://authorea.com/users/226952/articles/480342-scflow-a-scalable-and-reproducible-analysis-pipeline-for-single-cell-rna-sequencing-data

