## Supplementary Report Files 1 to 7 for "scFlow: A Scalable and Reproducible Analysis Pipeline for Single-Cell RNA Sequencing Data": File_1_Zhou_et_al_human_dimis_qc_report.html

scFlow - Quality Control Report


### Quality Control Report


### **scFlow** - Quality Control Report

###### 28 July, 2021

#### Overview

##### Post-QC Summary

- Cells  2,901
- Genes  14,371
- Total counts (M)  2931
- Total features (M)  1839
- Mitochondrial counts (M%)  3.45%
- Ribosomal counts (M%)  1.06%

\* M is the median for QC-passed cells

##### Key Parameters

- Library size  400 – 12,803
- Features/Genes  400 – 6,933
- Max. Mito. Counts  10.0%
- Ribo. Counts  0.0% – 100.0%
- Drop Mito. Genes  Yes
- Drop Ribo. Genes  No

##### Steps

Metadata Imported  ✔️    Ambient RNA / EmptyDrops Run  ❌    Thresholding  ✔️ ️   Gene/Cell QC Metric Annotation  ✔️ ️   Doublets/Multiplets Identified  (doubletfinder) ✔️

#### Sample metadata

A total of 18 metadata variables were imported from the sample sheet for this sample: -

#### Empty droplet identification

EmptyDrops was not run on this dataset.

#### Count depth distribution by barcode rank (high to low counts)

**Figure: Barcode count depth rank plot.** The ‘elbow’ indicates where count depth decreases rapidly (relative increase in background counts), and can be used to inform the count depth threshold. The applied lower-limit counts threshold is indicated at 400 counts (red line).

#### Number of counts / features per cellular barcode

The maximum number of counts per cell threshold was determined adaptively for this sample as >=4 median average deviations (MADs), or 12803 total counts per cell. The maximum number of features per cell threshold was determined adaptively for this sample as >=4 median average deviations (MADs), or 6933 total features per cell.

**Figure: Histogram of count depth per cell.** A lower-limit threshold of 400 and an upper-limit threshold of 12803 was applied (red line).

**Figure: Histogram of number of genes per cell.** A lower-limit threshold of 400 and an upper-limit threshold of 6933 was applied (red line).

#### Number of genes versus count depth

**Figure: Number of genes versus count depth coloured by relative mitochondrial counts.** The count-depth threshold of 400 – 12803 counts and the number of genes threshold of 400 – 6933 genes are indicated with vertical and horizontal red lines, respectively. Cells with high mitochondrial counts are typically in cells with relatively lower count depth. Cells with fractional mitochondrial counts higher than 0.100 (i.e. 10.00%) were filtered.

#### Fraction of mitochondrial / ribosomal counts

**Figure: Histogram of mitochondrial fraction per cell.** An upper-threshold of 0.100 (i.e. 10.00%) maximum mitochondrial fraction was applied (red line).

**Figure: Histogram of ribosomal fraction per cell.** An upper-threshold of 1 (i.e. 100.00%) maximum ribosomal fraction was applied (red line).

#### Doublet/multiplet identification

A total of 2901 cells which passed QC were submitted to the multiplet identification algorithm “doubletfinder”. This identified 67 multiplets and 2834 singlets. To identify these, the variable(s) “nCount\_RNA, pc\_mito” were first regressed out of the data. The first 10 principal components were used to identify the 2000 most variable genes. An assumed doublet formation rate of 0.023208 (i.e. 2.32% ) was applied.

##### Parameter sweep for optimal pK

A pK value of 0.005 was specified, therefore a parameter sweep was not performed.

##### Multiplet visualization in two-dimensional space

The 67 multiplets identified by “doubletfinder” are visualized below in red in PCA space, tSNE space, and UMAP space.

**PCA**

**tSNE**

**UMAP**

#### Full QC parameters and results

---

A report by **scFlow**
