## Supplementary Report Files 1 to 7 for "scFlow: A Scalable and Reproducible Analysis Pipeline for Single-Cell RNA Sequencing Data": File_2_Mathys_et_al_merged_report.html

scFlow - Merged Multi-Sample Quality-Control Report


### Merged Multi-Sample Quality-Control Report


### **scFlow** - Merged Multi-Sample Quality-Control Report

###### 29 July, 2021

#### Sample Pseudobulking Plots

##### Pseudobulked Sample Dimensionality Reduction Plots

These plots represent the pseudobulking of all cells from a sample (“*manifest*”) into a single point in two-dimensional space. Taken together with the merge summary plots, these may help to identify potential outliers or otherwise problematic samples. Data are visualized with: PCA, UMAP\_PCA.

##### PCA

##### UMAP\_PCA

##### Pseudobulked Sample Gene Expressivity Plot

Below, the pseudobulked counts matrix is used to determine which genes are detected or are absent in each sample. Hierarchical clustering with Euclidean distance is used to visualize the samples according to these binarized gene expressivity matrices. Samples with marked differences in the number and range of expressed genes may be potentially problematic and this facet of quality-control may be further explored in the related `total_features_by_counts` plots in this report.

**Figure: Heatmap and dendrogram of gene expressivity by sample.** Genes (rows) and samples (columns) are hierarchically clustered and reveal the number and distribution of expressed (purple) and non-expressed (yellow) genes per sample.

#### Cell Numbers Plot

The number of cells retained per sample (“*manifest*”) after quality-control. Samples which contribute relatively few cells to the study may require quality-control parameter optimization. In some cases, it may be preferable to drop such problematic samples from the study entirely.

**Figure: Number of cells per sample post-QC.**.

#### Multi-Sample Summary Plots

The multi-sample summary plots and matching data tables below examine key sample quality metrics (total\_features\_by\_counts, total\_counts, pc\_mito, pc\_ribo) across all experimental samples. These highlight samples with marked differences in these metrics and may be used to inform revision of quality-control thresholds. In more extreme cases (e.g. severe outliers, sample degradation, etc.), it may be appropriate to omit sample(s) from an analysis. To facilitate the identification of these potentially problematic samples, the *QC* column in the tables below highlight any samples with metrics falling outside of two or three standard deviations (σ) of the mean.

These metrics are also visualized across the specified facet variables:  *pathological\_diagnosis* .

#### total\_features\_by\_counts

**Figure: total\_features\_by\_counts by individual sample (manifest).**

**Table: total\_features\_by\_counts by individual sample (manifest).**

##### total\_features\_by\_counts\_vs\_pathological\_diagnosis

**Figure: total\_features\_by\_counts by pathological\_diagnosis.**

**Table: total\_features\_by\_counts by pathological\_diagnosis.**

---

#### total\_counts

**Figure: total\_counts by individual sample (manifest).**

**Table: total\_counts by individual sample (manifest).**

##### total\_counts\_vs\_pathological\_diagnosis

**Figure: total\_counts by pathological\_diagnosis.**

**Table: total\_counts by pathological\_diagnosis.**

---

#### pc\_mito

**Figure: pc\_mito by individual sample (manifest).**

**Table: pc\_mito by individual sample (manifest).**

##### pc\_mito\_vs\_pathological\_diagnosis

**Figure: pc\_mito by pathological\_diagnosis.**

**Table: pc\_mito by pathological\_diagnosis.**

---

#### pc\_ribo

**Figure: pc\_ribo by individual sample (manifest).**

**Table: pc\_ribo by individual sample (manifest).**

##### pc\_ribo\_vs\_pathological\_diagnosis

**Figure: pc\_ribo by pathological\_diagnosis.**

**Table: pc\_ribo by pathological\_diagnosis.**

---

scFlow v0.7.1 – 2021-07-29 11:34:00

---
