## Supplementary Report Files 1 to 7 for "scFlow: A Scalable and Reproducible Analysis Pipeline for Single-Cell RNA Sequencing Data": File_3_Ximerakis_et_al_integrate_report.html

scFlow - Integration Report


### Integration Report


### **scFlow** - Integration Report

###### 29 July, 2021

### Dataset Integration

Datasets were integrated using the method **‘Linked Inference of Genomic Experimental Relationships (LIGER)’** [@Welch2019]. This involved four pre-processing steps: (1) normalization for UMIs per cell, (2) subsetting the most variable genes for each dataset, (3) scaling by root-mean-square across cells, and (4) filtering of non-expressive genes.

**Key pre-processing parameters:**

- Number of variable genes per dataset (individual) selected for integration: **3000**
- Total number of variable genes used for integration (the union across all individuals): **4161**

**Note**: The length of the union across datasets (individuals) varies. The Venn and UpSet plots below may reveal outlying dataset/s.

**Upset chart of selected variable genes**: The first  ***16***  vertical bar charts show the sizes of isolated dataset participation to the total variable genes used for integration.

##### LIGER Factorization

An integrative non-negative matrix factorization was performed in order to identify shared and distinct metagenes (factors) across the datasets. The corresponding factor/metagene loadings were calculated for each cell.

**Key factorization parameters:**

- Number of factors (inner dimension of factorization; k): **20**
- Penalty parameter which limits the dataset-specific component of the factorization (lambda): **5**
- Resolution parameter which controls the number of communities detected: **1**

### Batch Effect Correction by LIGER

The performance of LIGER in batch effect correction was evaluated by comparison with dataset without a data integration algorithm applied (i.e. PCA input for dimensionality reduction). For each of the categorical covariates specified by the user two side-by-side comparisons have been represented: (1) visualisation of the batch effect using *tSNE* plots, and (2) quantification of the batch effect based on *kBET* test results [@Büttner2019].

In each *kBET* plot, the rejection rate represents the fraction of neighbourhoods with a label composition different from the global composition of batch labels. A significantly different observed vs. expected rejection rate opposes the well-mixedness of the data.

### Categorical covariates

#### manifest

**Figure: UMAP by manifest**

**Figure: UMAP (Liger) by manifest**

**Figure: kBET by manifest**

**Figure: kBET (Liger) by manifest**

#### group

**Figure: UMAP by group**

**Figure: UMAP (Liger) by group**

**Figure: kBET by group**

**Figure: kBET (Liger) by group**

### Clustering

Data generated by *PCA* and *LIGER* were used as inputs for clustering.

**Key clustering parameters:**

- Clustering method: **leiden**
- Number of nearest neighbors to use (k): **50**
- Resolution parameter that controls the resolution of clustering.: **0.001**

**Figure: UMAP**

**Figure: UMAP(Liger)**

scFlow v0.7.1 – 2021-07-29 17:12:32

---

A report by **scFlow**
