## Supplementary Report Files 1 to 7 for "scFlow: A Scalable and Reproducible Analysis Pipeline for Single-Cell RNA Sequencing Data": File_4_Mathys_et_al_celltype_metrics_report.html

scFlow - Cell-type and Cluster Overview Report


### Cell-type and Cluster Overview Report


### **scFlow** - Cell-type and Cluster Overview Report

###### 04 August, 2021

#### Dimensionality Reduction Plots

##### Cell-types

This two-dimensional embedding of cells should demonstrate spatial separation of major cell-types; if not, it may be useful to manually re-annotate cell-types with reference to cell-type associated marker genes, and/or revise dimensionality reduction and/or clustering parameters.

**Figure: UMAP\_Liger plot of cell-types.** Each cell is coloured by annotated cell-type.

###### Top Cell-type Marker Genes Table

###### Top Cell-Type Marker Genes Plot

**Figure: Top marker genes for cells grouped by cell-type.** Point colour represents expression and size represents the fraction of cells within the group expressing the gene.

##### Clusters

Optimal clustering parameters should provide a level of granularity sufficient to identify minority cell-types as distinct numbered clusters. If any cluster encompasses an adjacent small, well-separated cluster, the clustering parameters should be revised to allow for the annotation of the potential minority cell-type.

**Figure: UMAP\_Liger plot of clusters.** Each cell is coloured by annotated cluster.

###### Top Cluster Marker Genes Table

###### Top Cluster Marker Genes Plot

**Figure: Top marker genes for cells grouped by cluster.** Point colour represents expression and size represents the fraction of cells within the group expressing the gene.

##### Manifest

**Figure: UMAP\_Liger plot of cells by manifest.**

##### Pathological\_diagnosis

**Figure: UMAP\_Liger plot of cells by pathological\_diagnosis.**

##### Sex

**Figure: UMAP\_Liger plot of cells by sex.**

##### Individual

**Figure: UMAP\_Liger plot of cells by individual.**

#### Cell-type Proportion Plots

These plots illustrate the absolute cell numbers and relative proportions of cell-types in the overall experiment (“*all*”), and within the groups: *manifest, pathological\_diagnosis, sex, individual*. To examine statistically significant differences in cell-type proportions, see the Dirichlet model results and report(s).

##### All

**Figure: Absolute cell-type numbers by all.**

**Figure: Relative cell-type numbers by all.**

##### Manifest

**Figure: Absolute cell-type numbers by manifest.**

**Figure: Relative cell-type numbers by manifest.**

##### Pathological\_diagnosis

**Figure: Absolute cell-type numbers by pathological\_diagnosis.**

**Figure: Relative cell-type numbers by pathological\_diagnosis.**

##### Sex

**Figure: Absolute cell-type numbers by sex.**

**Figure: Relative cell-type numbers by sex.**

##### Individual

**Figure: Absolute cell-type numbers by individual.**

**Figure: Relative cell-type numbers by individual.**

#### Cell-type Metric Plots

These plots summarize the distribution of various cell metrics (*pc\_mito, pc\_ribo, total\_counts, total\_features\_by\_counts*) for each cell-type.

##### pc\_mito

**Figure: Average pc\_mito by cell-type**. Bars represent mean average +/- SEM.

**Figure: Distributions of pc\_mito by cell-type**.

##### pc\_ribo

**Figure: Average pc\_ribo by cell-type**. Bars represent mean average +/- SEM.

**Figure: Distributions of pc\_ribo by cell-type**.

##### total\_counts

**Figure: Average total\_counts by cell-type**. Bars represent mean average +/- SEM.

**Figure: Distributions of total\_counts by cell-type**.

##### total\_features\_by\_counts

**Figure: Average total\_features\_by\_counts by cell-type**. Bars represent mean average +/- SEM.

**Figure: Distributions of total\_features\_by\_counts by cell-type**.

scFlow v0.7.1 – 2021-08-04 14:59:15

---

A report by **scFlow**
