## Supplementary Report Files 1 to 7 for "scFlow: A Scalable and Reproducible Analysis Pipeline for Single-Cell RNA Sequencing Data": File_5_Mathys_et_al_Oligo_MASTZLM_Control_vs_pathological_diagnosisAD_de_report.html

scFlow - Differential Expression Analysis Report


### Differential Expression Analysis Report


### **scFlow** - Differential Expression Analysis Report

###### 09 August, 2021

##### Analysis summary

Differential gene expression was performed on a total of 1,864 genes which expressed a minimum of 1 count in at least 10 percent of the cells. Differentially expressed genes (DEGs) were detected at a fold-change threshold of 1.1 and an adjusted p-value (padj) cut-off of 0.05 was applied.

##### Result summary

**Up-regulated:** 3 (0.160944%)

**Down-regulated:** 25 (1.3412%)

##### Volcano plot

##### List of differentially expressed gene

scFlow v0.7.1 – 2021-08-09 10:19:00

---

A report by **scFlow**
