## Supplementary Report Files 1 to 7 for "scFlow: A Scalable and Reproducible Analysis Pipeline for Single-Cell RNA Sequencing Data": File_6_Mathys_et_al_Oligo_MASTZLM_Control_vs_pathological_diagnosisAD_DE_ipa_report.html

scFlow - Impacted Pathway Enrichment Report


### Impacted Pathway Enrichment Report


### **scFlow** - Impacted Pathway Enrichment Report

###### 09 August, 2021

### Pathway enrichment analysis

#### Enrichment analysis by WebGestaltR

##### Analysis summary

**Analysis name:** Oligo\_MASTZLM\_Control\_vs\_pathological\_diagnosisAD\_DE  
**Enrichment method for WebGestaltR:** ORA  
**Enrichment database:** geneontology\_Biological\_Process, pathway\_Reactome, pathway\_Wikipathway

##### Dot plots for top 10 enriched pathway

##### geneontology\_biological\_process

##### reactome

##### wikipathway

##### Result Table: geneontology\_biological\_process

##### Result Table: reactome

##### Result Table: wikipathway

#### Enrichment analysis by ROntoTools

Enrichment analysis using ROntoTools was not performed

#### Enrichment analysis by enrichR

##### Analysis summary

**Analysis name:** Oligo\_MASTZLM\_Control\_vs\_pathological\_diagnosisAD\_DE  
**Enrichment database:** GO\_Biological\_Process\_2018, Reactome\_2016, WikiPathways\_2019\_Human

##### Dot plots for top 10 enriched pathway

##### GO\_Biological\_Process\_2018

##### Reactome\_2016

##### WikiPathways\_2019\_Human

##### Result Table: GO\_Biological\_Process\_2018

##### Result Table: Reactome\_2016

##### Result Table: WikiPathways\_2019\_Human

scFlow v0.7.1 – 2021-08-09 10:39:04

---

A report by **scFlow**
