## Supplementary Report Files 1 to 7 for "scFlow: A Scalable and Reproducible Analysis Pipeline for Single-Cell RNA Sequencing Data": File_7_Ximerakis_et_al_dirichlet_report.html

scFlow - Cell-type Proportion Modeling Report


### Cell-type Proportion Modeling Report


### **scFlow** - Cell-type Proportion Modeling Report

###### 05 August, 2021

#### Analysis summary

Relative differences in the proportions of 14 different cell-types (*cluster\_celltype*) across 16 samples (*manifest*) in 2 experimental groups (*group:* Old, Young) were examined using a Dirichlet model. This model allows differences in cell-type composition to be tested while accounting for the proportions of all of the other cell-types.

Statistically significant differences in cell-type composition were observed in the following cell-types:  *TEINH13*

#### TEINH13 ✅

Statistically significant differences in the proportion of TEINH13 cells were observed between classes of *group* (vs Young) using a Dirichlet model.

**Figure: Relative proportion of TEINH13 by manifest.**

**Figure: Relative proportion of TEINH13 by group.** (Dirichlet-multinomial regression, `*`adjusted p ≤ 0.05, `**`adjusted p ≤ 0.01, `***`adjusted p ≤ 0.001); error bars: SEM.

##### Table: Dirichlet model p-values for TEINH13 (vs Young)

#### Oligo ❌

No statistically significant differences were observed in the proportion of Oligo cells between classes of *group* (vs Young) using a Dirichlet model.

**Figure: Relative proportion of Oligo by manifest.**

**Figure: Relative proportion of Oligo by group.** (Dirichlet-multinomial regression, `*`adjusted p ≤ 0.05, `**`adjusted p ≤ 0.01, `***`adjusted p ≤ 0.001); error bars: SEM.

##### Table: Dirichlet model p-values for Oligo (vs Young)

#### TEINH19 ❌

No statistically significant differences were observed in the proportion of TEINH19 cells between classes of *group* (vs Young) using a Dirichlet model.

**Figure: Relative proportion of TEINH19 by manifest.**

**Figure: Relative proportion of TEINH19 by group.** (Dirichlet-multinomial regression, `*`adjusted p ≤ 0.05, `**`adjusted p ≤ 0.01, `***`adjusted p ≤ 0.001); error bars: SEM.

##### Table: Dirichlet model p-values for TEINH19 (vs Young)

#### Endo ❌

No statistically significant differences were observed in the proportion of Endo cells between classes of *group* (vs Young) using a Dirichlet model.

**Figure: Relative proportion of Endo by manifest.**

**Figure: Relative proportion of Endo by group.** (Dirichlet-multinomial regression, `*`adjusted p ≤ 0.05, `**`adjusted p ≤ 0.01, `***`adjusted p ≤ 0.001); error bars: SEM.

##### Table: Dirichlet model p-values for Endo (vs Young)

#### Pericyte ❌

No statistically significant differences were observed in the proportion of Pericyte cells between classes of *group* (vs Young) using a Dirichlet model.

**Figure: Relative proportion of Pericyte by manifest.**

**Figure: Relative proportion of Pericyte by group.** (Dirichlet-multinomial regression, `*`adjusted p ≤ 0.05, `**`adjusted p ≤ 0.01, `***`adjusted p ≤ 0.001); error bars: SEM.

##### Table: Dirichlet model p-values for Pericyte (vs Young)

#### VSMC ❌

No statistically significant differences were observed in the proportion of VSMC cells between classes of *group* (vs Young) using a Dirichlet model.

**Figure: Relative proportion of VSMC by manifest.**

**Figure: Relative proportion of VSMC by group.** (Dirichlet-multinomial regression, `*`adjusted p ≤ 0.05, `**`adjusted p ≤ 0.01, `***`adjusted p ≤ 0.001); error bars: SEM.

##### Table: Dirichlet model p-values for VSMC (vs Young)

#### Ependymal ❌

No statistically significant differences were observed in the proportion of Ependymal cells between classes of *group* (vs Young) using a Dirichlet model.

**Figure: Relative proportion of Ependymal by manifest.**

**Figure: Relative proportion of Ependymal by group.** (Dirichlet-multinomial regression, `*`adjusted p ≤ 0.05, `**`adjusted p ≤ 0.01, `***`adjusted p ≤ 0.001); error bars: SEM.

##### Table: Dirichlet model p-values for Ependymal (vs Young)

#### TEGLU13 ❌

No statistically significant differences were observed in the proportion of TEGLU13 cells between classes of *group* (vs Young) using a Dirichlet model.

**Figure: Relative proportion of TEGLU13 by manifest.**

**Figure: Relative proportion of TEGLU13 by group.** (Dirichlet-multinomial regression, `*`adjusted p ≤ 0.05, `**`adjusted p ≤ 0.01, `***`adjusted p ≤ 0.001); error bars: SEM.

##### Table: Dirichlet model p-values for TEGLU13 (vs Young)

#### Micro ❌

No statistically significant differences were observed in the proportion of Micro cells between classes of *group* (vs Young) using a Dirichlet model.

**Figure: Relative proportion of Micro by manifest.**

**Figure: Relative proportion of Micro by group.** (Dirichlet-multinomial regression, `*`adjusted p ≤ 0.05, `**`adjusted p ≤ 0.01, `***`adjusted p ≤ 0.001); error bars: SEM.

##### Table: Dirichlet model p-values for Micro (vs Young)

#### OPC ❌

No statistically significant differences were observed in the proportion of OPC cells between classes of *group* (vs Young) using a Dirichlet model.

**Figure: Relative proportion of OPC by manifest.**

**Figure: Relative proportion of OPC by group.** (Dirichlet-multinomial regression, `*`adjusted p ≤ 0.05, `**`adjusted p ≤ 0.01, `***`adjusted p ≤ 0.001); error bars: SEM.

##### Table: Dirichlet model p-values for OPC (vs Young)

#### TEGLU10 ❌

No statistically significant differences were observed in the proportion of TEGLU10 cells between classes of *group* (vs Young) using a Dirichlet model.

**Figure: Relative proportion of TEGLU10 by manifest.**

**Figure: Relative proportion of TEGLU10 by group.** (Dirichlet-multinomial regression, `*`adjusted p ≤ 0.05, `**`adjusted p ≤ 0.01, `***`adjusted p ≤ 0.001); error bars: SEM.

##### Table: Dirichlet model p-values for TEGLU10 (vs Young)

#### VLMC ❌

No statistically significant differences were observed in the proportion of VLMC cells between classes of *group* (vs Young) using a Dirichlet model.

**Figure: Relative proportion of VLMC by manifest.**

**Figure: Relative proportion of VLMC by group.** (Dirichlet-multinomial regression, `*`adjusted p ≤ 0.05, `**`adjusted p ≤ 0.01, `***`adjusted p ≤ 0.001); error bars: SEM.

##### Table: Dirichlet model p-values for VLMC (vs Young)

#### PVM ❌

No statistically significant differences were observed in the proportion of PVM cells between classes of *group* (vs Young) using a Dirichlet model.

**Figure: Relative proportion of PVM by manifest.**

**Figure: Relative proportion of PVM by group.** (Dirichlet-multinomial regression, `*`adjusted p ≤ 0.05, `**`adjusted p ≤ 0.01, `***`adjusted p ≤ 0.001); error bars: SEM.

##### Table: Dirichlet model p-values for PVM (vs Young)

#### Astro ❌

No statistically significant differences were observed in the proportion of Astro cells between classes of *group* (vs Young) using a Dirichlet model.

**Figure: Relative proportion of Astro by manifest.**

**Figure: Relative proportion of Astro by group.** (Dirichlet-multinomial regression, `*`adjusted p ≤ 0.05, `**`adjusted p ≤ 0.01, `***`adjusted p ≤ 0.001); error bars: SEM.

##### Table: Dirichlet model p-values for Astro (vs Young)

#### Supplementary data tables

##### Table: Absolute cell numbers matrix

##### Table: Relative cell proportions matrix

##### Table: Normalized relative cell proportion table

Note: These are the final cell-type proportions and associated metadata used as inputs in the Dirichlet model. Differences in cell-type proportion values between this table and the previous relative proportions table may arise from the transformation of variables to cause the values to shrink away from extreme values of 0 and 1. See the *DirichletRegData* function in the DirichletReg package for more details.

scFlow v0.7.1 – 2021-08-05 14:47:39

---

A report by **scFlow**
